# Engineering the key central metabolic enzyme to utilize a non-canonical redox cofactor

**DOI:** 10.1101/2025.05.28.656713

**Authors:** Jin Young Kim, Edward King, Youtian Cui, Linyue Zhang, Justin B. Siegel, Han Li

## Abstract

Redesigning central metabolism enzymes to utilize non-canonical cofactors that are orthogonal to NAD^+^ and NADP^+^ may allow partitioning of reducing power into a distinct redox cofactor pool that is preserved for user-desired biosynthetic tasks, without being dissipated by the complex natural metabolism in the cells. Here, we engineer *Escherichia coli* glyceraldehyde-3-phosphate dehydrogenase (GapA), the key enzyme in glycolysis, to utilize nicotinamide mononucleotide (NMN^+^) while greatly reducing its original activity with its native cofactor, NAD^+^. Through rational design, we identify the variant A180S with a ∼3-fold increase in NMN^+^ specific activity compared to that of the wildtype (WT). Next, we design a double mutant, G10R-A180S (RS), where G10R mutation hampers the catalytic efficiency for NAD^+^ but also decreased the activity toward NMN^+^. Importantly, based on the hypothesis that the role of NAD^+^ in coordinating oligomer interface must be recapitulated in NMN^+^-utilizing enzymes, we develop a triple mutant G10R-A180S-G187Q (RSQ). Rosetta modeling predicts that the new mutation G187Q bridges separate subunits across the cofactor binding pocket, which possibly affords stronger association of the GapA tetramer, the active form of this enzyme. This final GapA variant features a ∼2.8 × 10^4^ switch of cofactor specificity from GapA’s natural cofactor NAD^+^ to the non-canonical cofactor NMN^+^. Overall, this work demonstrates the engineering of a highly conserved, central metabolic enzyme for the utilization of a non-canonical redox cofactor NMN^+^. The design principle discovered here may be broadly applicable to many enzymes that bind cofactors using an intersubunit pocket.

## Introduction

Metabolic pathways for biomanufacturing of fuels, chemicals, and commodities are often powered by the ubiquitous redox cofactor, nicotinamide adenine dinucleotide (phosphate) (NAD(P)H), which are mainly generated by the central metabolism such as Embden–Meyerhof–Parnas (EMP) glycolysis, pentose phosphate pathway (PPP), and Tricarboxylic Acid (TCA) cycle. However, NAD(P)H are also utilized by cellular respiration, fermentation, and biomass formation, leading to a loss of reducing power for the desired metabolic reactions. Thus, many attempts were reported to direct the reducing power to desired products by rewiring metabolic pathways of host cells^1–3^. To specifically direct reducing power and minimize electron dissipation to the side-reactions, recent efforts explored the non-canonical cofactors which operate in parallel to NAD(P)/H^4–11^. Nevertheless, the phosphorylating glyceraldehyde 3-phosphate dehydrogenase (GAPDH), one of the most active, conserved, and wide-spread enzymes in carbon and energy metabolism across all kingdoms of life^12^, has not been re-purposed to recycle non-canonical cofactors.

In this work, we specifically targeted the *Escherichia coli* GAPDH, which oxidizes glyceraldehyde-3-phosphate (G3P) to 1,3-bisphosphate-D-glycerate (1,3BPG) while reducing NAD^+^ to yield NADH. *E. coli* GAPDH (GapA) is particularly interesting to engineer due to its high catalytic turnover^13^ and high conservation of amino acid sequence^14^. These two factors are important as the high catalytic activity likely supports high turnover rate after protein engineering, and the high conservation of sequence likely leads to a translatable engineering strategy for GAPDHs from different organisms. Therefore, we envision the engineered *E. coli* GapA will serve as a tool to channel reducing equivalents into non-canonical cofactor pools with high efficiency by hijacking the universal EMP glycolysis that operates with high flux in divergent organisms.

Although there had been attempts to engineer GapA to be more active on NADP^+^,^15^ improving GapA’s activity for non-canonical cofactors has not been investigated. Among various non-canonical cofactors that mimic NAD(P)/H^16–22^, nicotinamide mononucleotide (NMN/H) is commonly targeted as a surrogate of the native NAD(P)/H due to its high structural similarity and low cost compared to the native cofactor^23^. Furthermore, it can be produced intracellularly through synthetic metabolic pathways in various organisms^24,25^, and has been shown to sustain central metabolism *in vivo*^8–10,26^ .Thus, we selected NMN/H as our non-canonical cofactor of interest for rational design and protein engineering.

Previously, we engineered the non-phosphorylating glyceraldehyde 3-phosphate dehydrogenase (GapN) from *Streptococcus mutans* to accept NMN^+^ and demonstrated its orthogonality *in vivo*^9^. Rosetta design^27^ was implemented to dock NAD^+^ and NMN^+^ in the GapN model to draw a rational design strategy that enabled improvement in NMN^+^ activity while reducing NAD^+^ activity. Although this was successful in that the resulting orthogonal GapN mutant (GapN Ortho) showed 465,000-fold increase in the catalytic efficiency compared to that of the wildtype, the mutations introduced in the GapN Ortho cannot be directly translated to GapA, because they belong to two structurally distinct enzyme classes: GapN belongs to the ‘aldehyde dehydrogenase family’, whereas GapA is classified as the ‘glyceraldehyde-3-phosphate dehydrogenase family’, according to the InterPro database^28^.

Nevertheless, our two-stage design strategy^11^ emerged from investigating a broad range of NMN(H)-dependent enzymes have again been proven effective in this work: First, we implemented Rosetta design and docking protocol to rationally select amino acid residues that are capable of interacting with the phosphate of NMN^+^. Next, novel interactions were designed to abolish the binding of the native cofactor NAD^+^ by closing the space in the conserved Rossmann-motif cavity.

Remarkably, this work discovered that connecting two halves of the cofactor binding pocket that belong to different subunits of GapA with a continuous hydrogen bond network further enhanced catalytic efficiency toward NMN^+^, without compromising orthogonality against natural cofactor NAD^+^. Besides its catalytic role, NAD^+^ can often play a structural role to bridge different subunits, when its binding pocket is situated at or near an intersubunit cleft. Non-canonical cofactors may not recapitulate this structural role, which has been largely overlooked in previous studies. Since a vast majority of cofactor-dependent enzymes exist as oligomers^29^ which may provide fitness gain in evolution^30^, this work reveals an imperative new aspect in non-canonical cofactor enzyme design that may be generalized to other redox enzymes.

## Results and Discussion

### Improving NMN^+^ binding of GapA

Structurally, NMN/H differs from NAD(P)/H that it is missing the adenosine monophosphate (AMP) tail while sharing the nicotinamide ring that is necessary for hydride transfer during redox catalysis (Figure 1A). Consistently, *K*_M_ for NMN^+^ is much poorer for GapA WT compared to NAD^+^ (Table 1). To improve NMN^+^ binding, we hypothesized that introducing more polar residues in the first shell of the cofactor binding site would be beneficial as we were able to apply this strategy to various enzymes previously^8,9,31^. Novel polar contacts should be established around the phosphate group or the ribose hydroxyl group as NMN^+^ is missing its AMP moiety (Figure 1A). Although there is a dimeric crystal structure solved for *E. coli* GapA (PDB: 1GAD), the active form of GAPDH is a tetramer. Thus, we structurally aligned the dimeric 1GAD to a homologous GapA from *Oryctolagus cuniculus* (*O. cuniculus*) (PDB: 1J0X) to build a tetrameric model that completes the cofactor binding site. We docked a conformer library of NMN^+^ into our tetrameric model to find the optimal binding pose of NMN^+^ in the complete binding site using Rosetta design and dock. Upon investigation of the complete cofactor binding site with NMN^+^, we selected five candidate residues that are capable of forming polar contacts with NMN^+^ when mutated to polar amino acids (Figure 1B). V185 is in the interfacial loop that could form novel intermolecular interactions, which would not have been identified if only the monomeric structure was investigated. G10 sits at the tip of the Rossmann alpha helix near the phosphate group of NMN^+^ as A180 also appears near the phosphate group but located on a loop running across from the Rossmann alpha helix. L100 and G120 are located on the loop running over the ribose and in the proximity of the hydroxyl groups of NMN^+^.

**Table 1.** Apparent kinetic parameters for *E. coli* GapA mutants.

| Enzyme | Cofactor | $k_{cat}$ (s<sup>-1</sup>) | $K_M$ (mM) | $k_{cat}/K_M$ (mM<sup>-1</sup> s<sup>-1</sup>) |
| --- | --- | --- | --- | --- |
| WT | NAD<sup>+</sup> | $6.3 \pm 0.2$ | $0.13 \pm 0.01$ | $48.4 \pm 2.7$ |
| | NADP<sup>+</sup> | $(5.2 \pm 0.1) \times 10^{-2}$ | $3.3 \pm 0.2$ | $(1.6 \pm 0.1) \times 10^{-2}$ |
| | NMN<sup>+</sup> | $(2.5 \pm 0.2) \times 10^{-2}$ | $18.0 \pm 2.3$ | $(1.4 \pm 0.1) \times 10^{-3}$ |
| A180S | NAD<sup>+</sup> | $2.1 \pm 0.4$ | $0.10 \pm 0.04$ | $23.5 \pm 7.9$ |
| | NADP<sup>+</sup> | $(1.5 \pm 0.3) \times 10^{-2}$ | $2.2 \pm 0.9$ | $(6.9 \pm 1.5) \times 10^{-3}$ |
| | NMN<sup>+</sup> | $(2.9 \pm 0.2) \times 10^{-2}$ | $8.1 \pm 1.4$ | $(3.7 \pm 0.5) \times 10^{-3}$ |
| G10R-A180S (RS) | NAD<sup>+</sup> | $(3.1 \pm 0.4) \times 10^{-2}$ | $5.5 \pm 0.6$ | $(5.7 \pm 1.3) \times 10^{-3}$ |
| | NADP<sup>+</sup> | $(8.5 \pm 0.8) \times 10^{-3}$ | $2.4 \pm 0.5$ | $(3.5 \pm 0.4) \times 10^{-3}$ |
| | NMN<sup>+</sup> | $(1.5 \pm 0.2) \times 10^{-2}$ | $10.4 \pm 2.1$ | $(1.5 \pm 0.2) \times 10^{-3}$ |
| G10R-A180S-G187Q (RSQ) | NAD<sup>+</sup> | $(4.1 \pm 0.4) \times 10^{-2}$ | $7.6 \pm 0.4$ | $(5.3 \pm 0.8) \times 10^{-3}$ |
| | NADP<sup>+</sup> | $(7.3 \pm 1.3) \times 10^{-3}$ | $1.8 \pm 0.7$ | $(4.4 \pm 1.4) \times 10^{-3}$ |
| | NMN<sup>+</sup> | $(3.4 \pm 0.1) \times 10^{-2}$ | $7.7 \pm 0.3$ | $(4.4 \pm 0.1) \times 10^{-3}$ |

**Figure 1.**
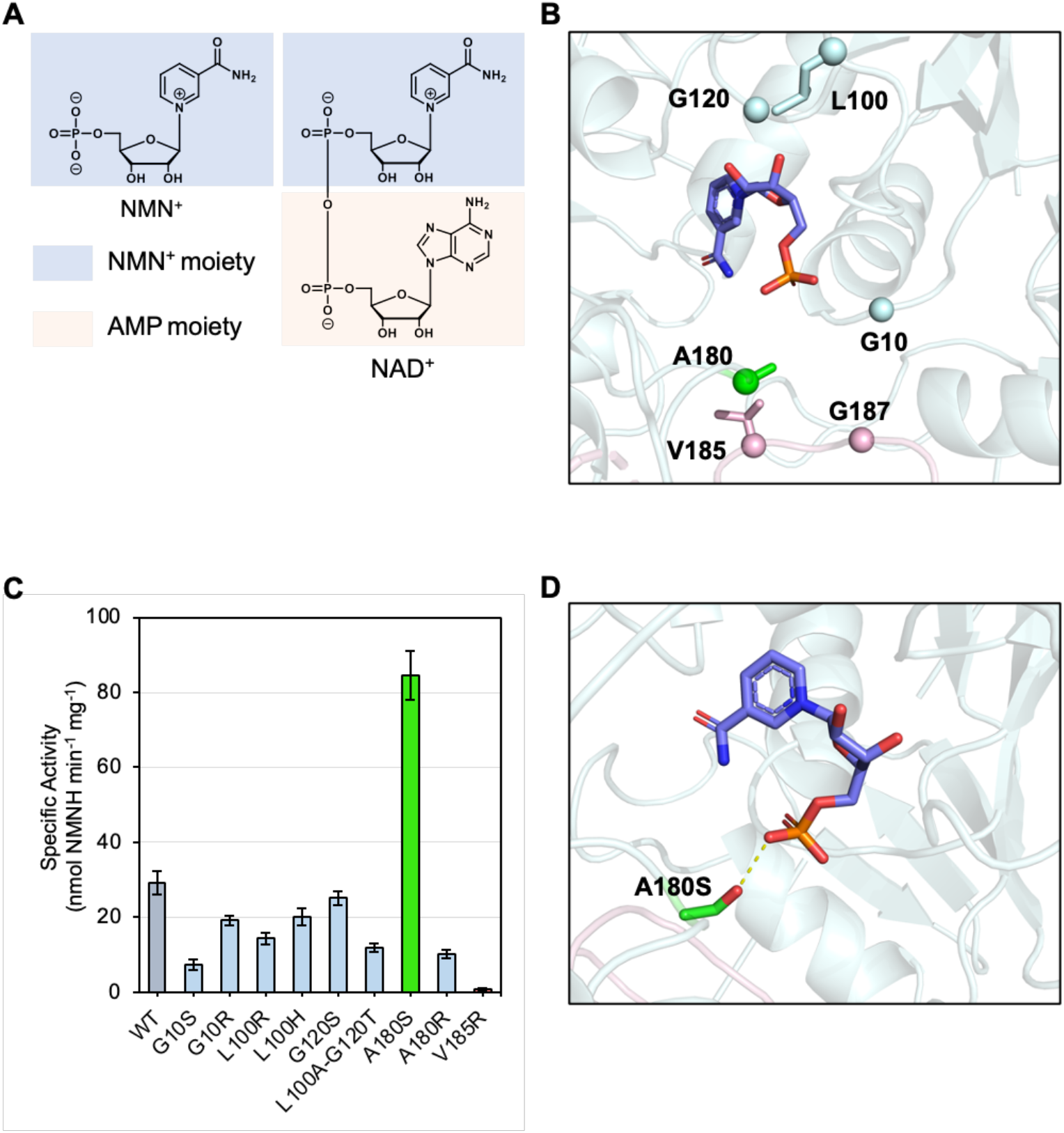
Engineering GapA WT for improving NMN^+^ activity. (A) Structures of NAD^+^ and NMN^+^. Structurally, NMN^+^ differs from NAD^+^ by the absence of the AMP moiety. (B) Model of GapA WT docked with NMN^+^. The cofactor is shown in slate. Each subunit is colored differently (e.g., the chains in lightblue and pink represent distinct subunits). The residue that increased NMN^+^ activity is highlighted in green. (C) Specific activity of rationally designed mutations for increasing NMN^+^ activity. Error bars represent one standard deviation of three replicates. (D) Predicted polar interaction between A180S and NMN^+^. A novel polar interaction between the phosphate group of NMN^+^ and the side chain of A180S is shown as a dashed line.

We built polar mutations on those five candidate residues to introduce polar contacts at the phosphate or hydroxyl group of NMN^+^. Serine and arginine were chosen as the polar mutations for G10 and A180, since each represents short and long polar amino acids respectively, and there are few successful examples reported with this approach^8,9,31^. For V185, only arginine was tested since V185 is located on the distant loop of another subunit. Thus, short polar amino acids would not be sufficient to form polar contacts. The distances from Cα atoms of G10 and A180 to the phosphate group of NMN^+^ are 5.5 and 4.8 Å respectively; however, the distance for V185 is 10.6 Å, which is twice as far as that of the other two residues (Figure S1). For L100 and G120, we not only designed mutations that could form polar interactions but also introduced mutations that could form hydrogen bonds with the hydroxyl groups of ribose. As a result, we tested nine mutations as an initial round of rational design screening (Figure 1C). We found a positive hit in the first round, A180S, which exhibited a specific activity of 84.5 ± 6.5 nmol min^-1^ mg^-1^ with NMN^+^, a 2.9-fold increase compared to the WT’s specific activity of 29.3 ± 3.1 nmol min^- 1^ mg^-1^. This first round of rational design was conducted by expressing and purifying the variants in *E. coli* BL21, a classic strain optimized for protein expression. However, this strain still harbors GapA WT which may form heterotetramer with the GapA variants and be co-purified. This convolutes the kinetic parameter determination. For better characterization of A180S, we constructed a W3CG strain with a *pncC* knock out (See Materials and Methods). This strain has its endogenous GapA disrupted^32^, and the deletion of *pncC* prevents degradation of NMN^+^. A180S was transformed into our newly constructed W3CG Δ*pncC* strain and assayed for its kinetic parameters. The kinetic parameters were measured to be *k*_cat_ (1.4 ± 0.1) x 10^−2^ s^-1^, *K*_m_ 5.1 ± 0.5 mM^-1^, and *k*_cat_/*K*_m_ (2.7 ± 0.2) x 10^−3^ mM^-1^ s^-1^ (Table 1). *K*_m_ was reduced by 3.5-fold compared to that of the WT (18.0 ± 2.3 mM^-1^), leading to an increase in the catalytic efficiency.

A Rosetta model was built for A180S with NMN^+^ to elucidate the mechanism of activity increase for NMN^+^ (Figure 1D). The model showed that A180S forms a hydrogen bond with the phosphate group of NMN^+^ at a distance of 2.8 Å as we hypothesized. This novel interaction reinforces the binding of NMN^+^ by anchoring its phosphate in the binding site in a catalytically active orientation. Although A180S showed a significant increase in NMN^+^ activity, its activity for the native cofactor NAD^+^ was still considerably high (Figure 2A, Table 1). For potential application as an *in vivo* catalyst, the promiscuous activity of this enzyme toward NAD^+^ will hamper efficient partitioning of reducing power from glycolysis to the non-canonical cofactor pool. Therefore, we next sought to decrease A180S’s activity toward NAD^+^.

**Figure 2.**
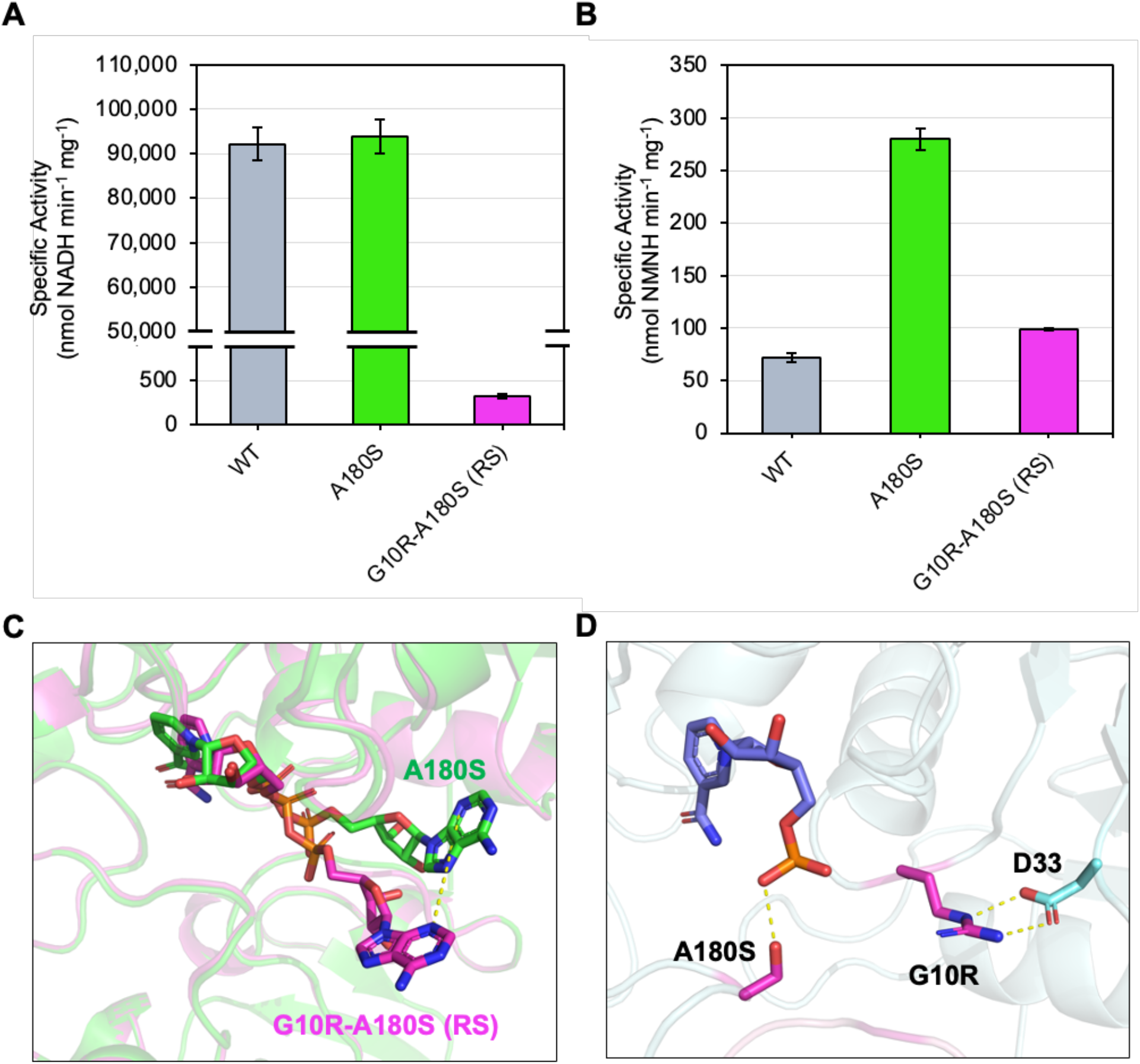
Increasing orthogonality of GapA A180S. (A) Specific activity of WT, A180S, and G10R-A180S (RS) toward the native cofactor, NAD^+^. Error bars represent one standard deviation of three replicates. (B) Specific activity of WT, A180S, and G10R-A180S (RS) toward the non-canonical cofactor, NMN^+^. Error bars represent one standard deviation of three replicates. (C) Superimposed Rosetta models of A180S and G10R-A180S (RS) with NAD^+^. The AMP moiety of NAD^+^ shifts outward from the binding pocket in the RS model due to the G10R mutation. A180S is shown in green, while RS is shown in magenta. Both models were generated using identical Rosetta constraints for NAD^+^ docking. (D) Predicted molecular interactions in the orthogonal G10R-A180S (RS) mutant. NMN^+^ is shown in slate, mutations in magenta, and the native amino acid residues in lightblue. Novel polar interactions between the side chains of mutations and NMN^+^ or the nearby residue, D33, are shown in dashed lines.

### Improving orthogonality by disrupting GapA’s native cofactor binding

To design mutations that could block binding of the native cofactor while minimally disturbing NMN^+^ binding, we applied a design principle of introducing steric hindrance to the AMP moiety of NAD^+^, which is not present in NMN^+^. The AMP group fits into a cleft formed between the alpha helix and second beta strand of the Rossmann fold. Occupying this space would occlude the adenosine from fitting, while the NMN^+^ binding pose would be nominally affected as NMN^+^ makes no interaction in that region.

Based on this principle, we designed G10R-A180S (RS). This double mutant showed drastically lower specific activity for NAD^+^ (Figure 2A). Its activity was measured to be 321 ± 20.4 nmol min^-1^ mg^-1^, a ∼290-fold decrease compared to that of WT and A180S ((9.22 ± 0.4) x 10^4^ nmol min^-1^ mg^-1^ and (9.39 ± 0.4) x 10^4^ nmol min^-1^ mg^-1^ respectively). Kinetic parameter characterization also showed that RS features a significant increase in *K*_m_ by 42-fold and decrease in *k*_cat_/*K*_m_ by 8491-fold for NAD^+^ compared to those of the WT (Table 1). However, RS also showed a decrease in activity toward NMN^+^ compared to A180S (Figure 2B, Table 1) which necessitates further engineering as described below.

To illustrate the mechanism by which a single mutation G10R abolish NAD^+^ activity by several thousand-fold, we conducted Rosetta modeling for RS with NAD^+^ docked. Compared to the A180S model, NAD^+^ in the RS model displayed an outward shift of the AMP moiety to the solvent, resulting in an unfavorable cofactor binding mode (Figure 2C). The distance of cofactor shift was as far as 7 Å, when the distance between the N3 atoms of adenine in A180S and RS was considered. This shift in cofactor tail is predicted to be caused by the occupation of binding site by a salt bridge formed between G10R and D33 (Figure 2D). The strategy of sealing the AMP cleft with electrostatic interaction pairs has been demonstrated before to result in NMN-orthogonal enzymes^11^.

The RS model docked with NMN^+^ predicted that G10R may not interfere with the binding of NMN^+^ (Figure 2D). To the contrary, both *K*_M_ and *k*_cat_ for NMN^+^ worsened for RS compared to A180S (Figure 2B, Table 1). This suggest that other factors that govern the enzyme activity, which were not captured in the static Rosetta model, need to be considered to rescue the NMN^+^ activity.

### Rejuvenating NMN^+^ activity while maintaining orthogonality

We hypothesized that building a stronger intermolecular interaction between subunits and further supporting of the void caused by missing AMP moiety would enhance NMN^+^ activity. Previous study has shown that GapA tetramer successively binds four NAD^+^ molecules one at a time, with one NAD^+^ binding to each subunit increasing the affinity of the next NAD^+^. This cooperative behavior in GapA enzyme activity may be mediated by better organized subunit interfaces upon cofactor binding, because NAD^+^ binding has been shown to reduce the excessive structural dynamic of the GapA^33^. While NAD^+^ can serve as cement between different subunits thanks to the extensive interactions it makes with both monomers lining the active site, NMN^+^ is limited in this ability due to fewer contacts with the binding pocket.

G187 was selected as the target residue as it is located in the loop that sits at the interface of two subunits and running close to the cofactor binding pocket (Figure 1B). We constructed six triple mutants based on the RS mutant as the parent, with G187 substituted to A, H, L, M, N, and Q. Two possibilities of introducing intermolecular interactions and filling the void were investigated: one by hydrophobic interaction with the nearby M43 in the interfacial helix and the other by polar interaction with the NMN^+^ phosphate or G10R (Figure S2). We selected A, H, L, M, N, and Q to examine the effect of different length and size for both hydrophobic and polar amino acids.

Interestingly, RS-G187Q (RSQ) showed a boost in NMN^+^ activity compared to RS, whereas other five mutants did not or had limited improvement (Figure 3A). The specific activity of RSQ for NMN^+^ was measured to be 268.3 ± 10.2 nmol min^-1^ mg^-1^, which is a 2.7-fold increase from RS and reached similar level to that of A180S. All six triple mutants also showed lower activity for both NAD^+^ and NADP^+^ (Figure 3B and 3C) than the RS, presumably because the mutations on G187 also shift the AMP moiety of canonical cofactors further out of binding pocket. Since RSQ had the highest specific activity for NMN^+^, we further characterized RSQ for kinetic parameters and Rosetta modeling.

**Figure 3.**
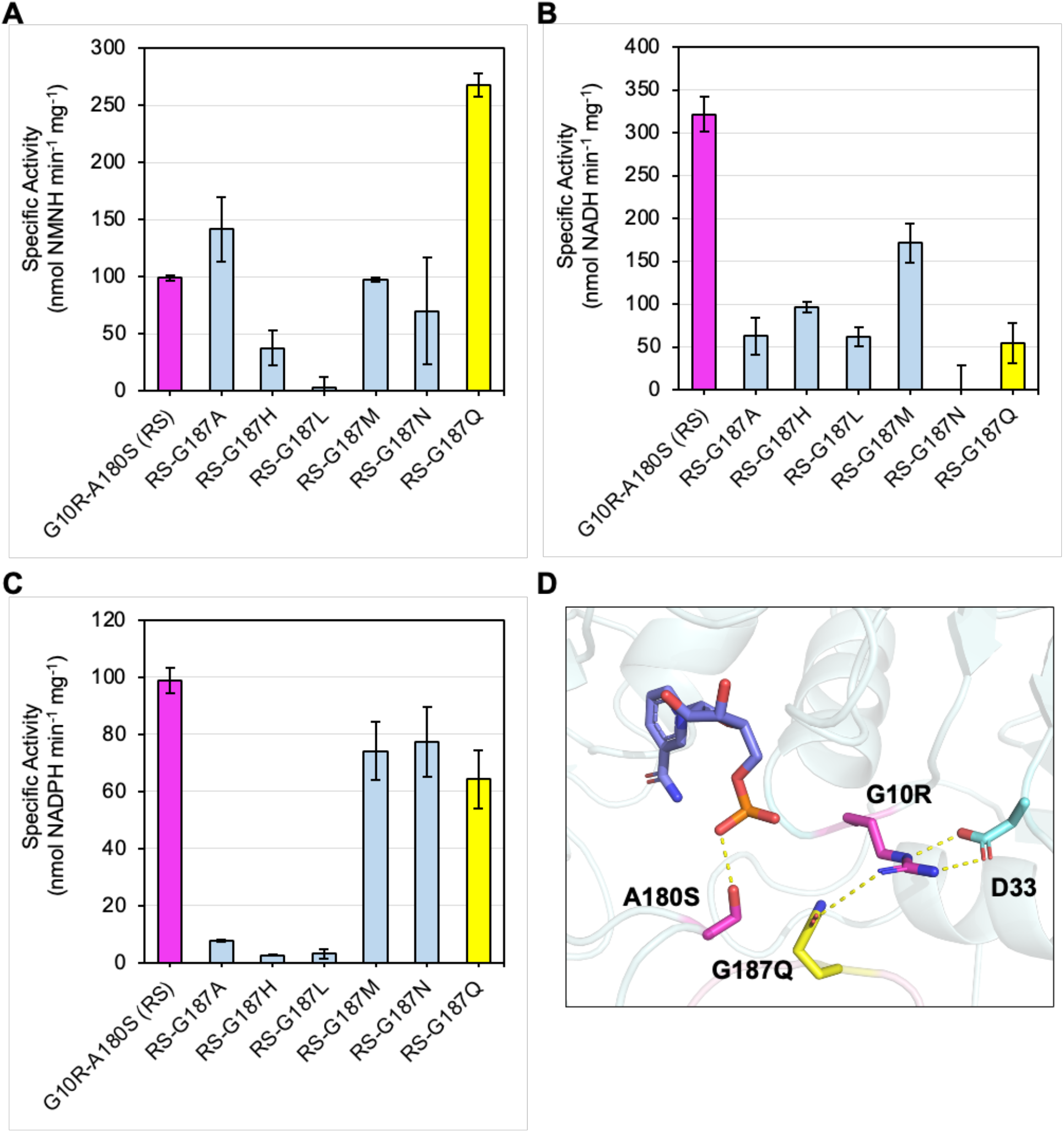
Improving NMN^+^ activity of the orthogonal mutant GapA G10R-A180S (RS). (A) Specific activity of RS and its second-round rationally designed variants toward NMN^+^. (B) Specific activity of the same variants toward NAD^+^. (C) Specific activity of the same variants toward NADP^+^. (D) Predicted molecular interactions of the mutant G10R-A180S-G187Q (RSQ). NMN^+^ is shown in slate, parental mutations in magenta, and the additional G187Q mutation in yellow. Novel polar interactions between the side chains of mutations and NMN^+^ or the nearby residue, D33, are shown in dashed lines. Error bars represent one standard deviation of three replicates.

The kinetic parameters of RSQ for NMN^+^ indicated a revitalization of NMN^+^ activity from RS by G187Q mutation (Table 1). The catalytic efficiency of RSQ was increased by 2.9-fold from that of the parent RS and became comparable to that of A180S. While invigorating the non-canonical cofactor activity, RSQ still maintained its orthogonality toward both NAD^+^ and NADP^+^ in a degree comparable to RS. In particular, the *K*_M_ for NAD^+^ and NADP^+^ of RSQ reached 7.6 mM and 1.8 mM respectively (Table 1), well above their cofactors’ physiological concentrations *in vivo*. This suggests that RSQ would be a suitable tool to establish NMN-regenerating EMP glycolysis inside the cells.

We built a Rosetta model of RSQ with NMN^+^ to further investigate the mechanism of improvement in NMN^+^ activity on top of RS. Our model showed that G187Q is forming a polar interaction with G10R (Figure 3D). G187Q, which belongs to a different subunit than G10R and D33, is predicted to form a continuous, intersubunit hydrogen bonding network with these two residues. This hydrogen bonding network may stabilize the enzyme’s oligomeric state and mitigate counter-productive flexibility within the cofactor binding pocket^26^. Furthermore, G187Q is positioned to add more steric hindrance to block NAD(P)^+^ binding.

## Conclusion

Manipulating redox driving force with a non-canonical cofactor is challenging due to an enzyme’s high affinity toward its native cofactor, especially if the enzyme is involved in the key central metabolism such as glycolysis. Here, we implemented a rational design workflow to search for potential mutations that would benefit NMN^+^ binding while maintaining orthogonality by hindering NAD^+^ binding of *E. coli* GapA. Our best performing variant, G10R-A180S-G187Q (RSQ) improved the catalytic efficiency for NMN^+^ by ∼3.1-fold while severely diminishing the catalytic efficiency for NAD^+^ by ∼9132-fold and NADP^+^ by ∼363-fold compared to the WT. Overall, this translates to ∼2.8 × 10^4^ switch of cofactor specificity from GapA’s natural cofactor NAD^+^ to the non-canonical cofactor NMN^+^. Structural investigation with Rosetta design and docking illustrated that blocking of the AMP tail by a bulky amino acid, filling the void of binding pocket caused by missing AMP tail, and bolstering intermolecular interactions between subunits are effective in not only improving non-canonical cofactor activity but also reducing canonical cofactor activity for GapA. Future work is needed to evaluate the *in vivo* utility of the engineered GapA to allocate reducing power in the form of NMNH, which is protected from consumption by the cells natural enzymes and can be specifically designated to desired synthetic pathways.

## Materials and Methods

### Plasmid and strain construction

The plasmids and strains used in this study are listed in Table S1. Plasmid construction was performed using the Gibson isothermal DNA assembly method^34^. The *E. coli gapA* gene was amplified from *E. coli* BW25113 genomic DNA and cloned into the pQE vector backbone (N-terminal 6x His-tag, ColE1 *ori*, Amp^R^) to generate the expression plasmid pEK-28. The W3CG Δ*pncC* strain was constructed by deleting the *pncC* gene in W3CG via P1 phage transduction using Keio collection, followed by flippase recombinase-mediated excision of the corresponding kanamycin resistance cassette^35^.

Site-directed mutagenesis was performed via PCR with KOD One DNA polymerase (Toyobo) and mutagenic primers carrying the target codon substitutions. pEK-28 was used as the template for single, double, or triple mutations constructed in this study. Two fragments of DNA, one carrying desirable mutations and the other one carrying the backbone of the plasmid, were ligated using the Gibson assembly method^34^. Gibson products were transformed into *E. coli* XLI-Blue (Stratagene) for cloning. Successfully cloned plasmids were transformed into either *E. coli* BL21(DE3) (Invitrogen) or W3CG Δ*pncC* for protein expression.

### Protein expression and purification

BL21(DE3) or W3CG Δ*pncC* cells harboring the plasmids encoding WT *gapA* or its variants were inoculated into 2xYT medium (Fisher Scientific) with 100 µg/mL of ampicillin for overnight culture at 37 °C. A 1% inoculum of the overnight culture was used to seed 50 mL of fresh 2xYT medium with 100 µg/mL of ampicillin until OD600 reached ∼0.6. The cultures were induced with 0.5 mM isopropyl β - D-1-thiogalactopyranoside (IPTG; Zymo Research Corporation) and incubated overnight at 30 °C with shaking at 250 rpm. Pelleted cells were resuspended in 50 mM Tris-Cl pH 7.7 buffer with 300 mM NaCl and 10 mM imidazole and disrupted with 0.1mm glass beads (BioSpec). Cell lysates were collected after centrifugation at 15000 rpm, 4 °C. Buffers used for the protein purification were modified from the His-Spin Protein Miniprep kit (Zymo Research Corporation) by switching their buffer salts from sodium phosphate to Tris-Cl. Cell lysates were purified with Ni-NTA resin (Thermo Scientific) and the modified buffers according to the kit manufacturer’s protocol. Protein concentrations were determined using the Bradford assay.

### GapA enzymatic assay and kinetic study

GapA specific activity was measured by adapting the assay protocol from the previous work^9^. Reactions were initiated by adding purified enzymes to an assay mixture containing 50 mM Tris-Cl pH 8.5, 0.2 mM EDTA, 50 mM Na_2_PO_4_, 3 mM DL-glyceraldehyde 3-phosphate (DL-G3P), and 5 mM cofactor at 25°C. Cofactor reduction was monitored with spectrophotometer (Molecular Devices) at 340 nm. All specific activities were normalized by subtracting no-substrate controls and the concentrations of enzymes used in the reaction.

Determination of the Michaelis-Menten kinetic parameters was completed by varying the concentration of cofactors in the same reaction mixture used for the specific activity assay. The enzyme concentration was set to either 0.2 or 2 μM depending on the activity of the enzymes toward the cofactor they were measured with. Initial reaction rates were recorded and fit to the Michaelis-Menten equation where *v*_0_ is the initial velocity, *E*_t_ is the total enzyme concentration, and *S* is the cofactor concentration.

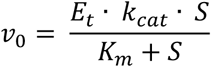

### Rosetta ligand docking and design

Due to the absence of a crystal structure of *E. coli* GapA in its active tetrameric state with a bound native cofactor, a homology model was built to represent the active tetramer of *E. coli* GapA. The dimeric structure of *E*.*coli* GapA (PDB: 1GAD) was structurally aligned to each subunit of the homologous rabbit muscle (*O. cuniculus*) GapA (PDB: 1J0X), which is resolved in an active tetrameric form with NAD^+^. The resulting model recapitulated the tetrameric symmetry and the active binding mode of NAD^+^, enabling formation of intersubunit polar contacts across the active sites. All subsequent simulations were based on this model.

Conformer libraries for NAD^+^ and NMN^+^ were constructed and optimized to preserve catalytically active geometries, following the protocol established in our previous study^9^. The Rosetta design protocol was implemented to design rotamers of amino acid substitutions predicted to improve NMN^+^ binding affinity or to evaluate model outputs on NAD^+^ binding. Each design cycle included multiple rounds of Monte Carlo sampling involving rigid body translation and rotation of ligand conformers, optimization of side chain hydrophobic and polar interactions, and backbone minimization to relieve steric hindrance around the binding site. Key interactions from the catalytically competent geometry for hydride transfer were constrained by manually setting distances and angles adopted from catalytically active conformations.

Mutations were introduced using the MutateResidue mover in RosettaScripts. For each design round, 500 models were generated, and the top 20 models were selected based on the protein-ligand interface energy and the Rosetta total energy score. These selected models were further manually inspected to choose the best model for each round of design. The codes used for RosettaScripts XML files, ligand parameters files, and constraint files used in this study are provided in the Supporting Information.

## Author Contribution

H.L., J.Y.K., E.K. designed the experiments. H.L., J.Y.K., E.K., and L.Z. performed the experiments and analyzed the results. J.Y.K., Y.C., and J.B.S. designed and performed Rosetta modeling. All authors wrote the manuscript.

## Notes

The authors declare no competing financial interest.

## Acknowledgment

H.L., J.Y.K., E.K., and L.Z. acknowledge support from the National Science Foundation (NSF) (Award no. 2328145), the National Institutes of Health (NIH) (Award no. 1R35GM153401-01), and Sloan Research Fellowship. Y.C. and J.B.S. acknowledge the funding of the National Institute of Environmental Health Sciences (Grant no. P42ES004699), the NIH (Grant no. R01 GM 076324-11), and the NSF (Grant nos. 1627539, 1805510, and 1827246).

## Notes

### Competing Interest Statement

The authors have declared no competing interest.

