## Supporting Information for "Engineering the key central metabolic enzyme to utilize a non-canonical redox cofactor"

**Table of Contents**

**A. Supplemental Figures and Tables**

Figure S1. Rosetta model of GapA docked with NMN+

Figure S2. Structural depiction of interfacial residue G10, M43, and G187

Table S1. Plasmids and strains used in this study

**B. Supplementary Text**

Example of Rosetta design and docking commands

**C. References**

**A. Supplemental Figures and Tables**

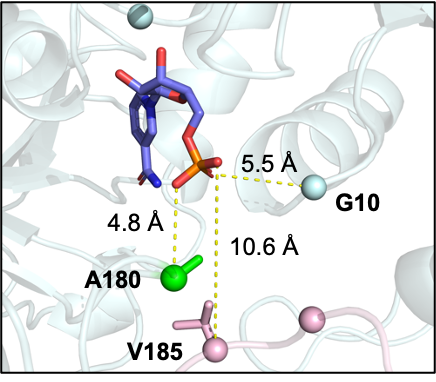

**Figure S1.** A model of *E. coli* GapA WT docked with NMN^+^. Distance from the phosphate group of NMN^+^ and the C_α_ of G10, A180, and V185 are visualized.

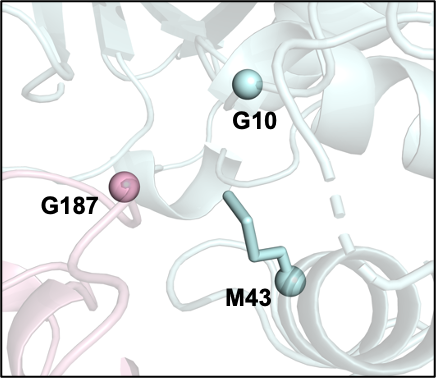

**Figure S2.** A model of GapA WT depicting the interface region of G187, G10, and M43. Each chain is represented in different colors.

**Table S1. Plasmids and strains used in this study**

| **Strains** | **Description** | **Reference** |
| --- | --- | --- |
| XLI-Blue | Cloning strain | Stratagene |
| BL21 (DE3) | Initial protein expression strain | Invitrogen |
| BW25113 | *lacI*^q^ *rrnB*_T14_ Δ*lacZ*_WJ16_ *hsdR514* Δ*araBAD*_AH33_ Δ*rhaBAD*_LD78_ | Datsenko et al.^1^ |
| W3CG | *E. coli* F-, *λ^-^*, *gapA10::Tn10*, *IN(rrnD-rrnE)1*, *rph-1* | Valverde et al.^2^ |
| **Plasmids** | **Description** | **Reference** |
| pQE | N-terminal 6x His-tag; ColE1 *ori*; Amp^R^; *P*_LlacO1_ | Li et al.^3^ |
| pEK-28 | pQE *E. coli* *gapA* | This study |
| pEK-30 | pQE *E. coli* *gapA* G10S | This study |
| pEK-31 | pQE *E. coli* *gapA* G10R | This study |
| pEK-38 | pQE *E. coli* *gapA* L100R | This study |
| pEK-39 | pQE *E. coli* *gapA* L100H | This study |
| pEK-36 | pQE *E. coli* *gapA* G120S | This study |
| pEK-37 | pQE *E. coli* *gapA* L100A-G120T | This study |
| pEK-32 | pQE *E. coli* *gapA* A180S | This study |
| pEK-33 | pQE *E. coli* *gapA* A180R | This study |
| pEK-35 | pQE *E. coli* *gapA* V185R | This study |
| pEK-52 | pQE *E. coli* *gapA* G10R-A180S | This study |
| pJK-17 | pQE *E. coli* *gapA* G10R-A180S-G187A | This study |
| pJK-18 | pQE *E. coli* *gapA* G10R-A180S-G187H | This study |
| pJK-19 | pQE *E. coli* *gapA* G10R-A180S-G187L | This study |
| pJK-20 | pQE *E. coli* *gapA* G10R-A180S-G187M | This study |
| pJK-21 | pQE *E. coli* *gapA* G10R-A180S-G187N | This study |
| pJK-22 | pQE *E. coli* *gapA* G10R-A180S-G187Q | This study |

**B. Supplementary Text**

Command:

/share/siegellab/kschu/software/Rosetta/main/source/bin/rosetta_scripts.default.linuxgccrelease -database /share/siegellab/software/kschu/Rosetta/main/database @GapA_NMN_flags -user_tag $SLURM_ARRAY_TASK_ID -out:suffix $SLURM_ARRAY_TASK_ID

Example of a flags file:

####UNIQUE TO RUN####

-nstruct 10

-parser:protocol GapA_3_mut.xml

###MUTATION SETUP###

-parser:script_vars target1=340P new_res1=ARG

-parser:script_vars target2=510P new_res2=SER

-parser:script_vars target3=187P new_res3=GLN

####SHARED FILES###

-enzdes::cstfile NMN.enzdes.cst

-in:file:s A180S_G10R_NMN.pdb

-packing::unboundrot A180S_G10R_NMN.pdb

-extra_res_fa NMN.params

-out:path:all /share/siegellab/jinyohg/docking/Final/GapA/NMN/Result

####SYSTEM SETUP###

-run::preserve_header

-run::version

-nblist_autoupdate

-linmem_ig 10

-jd2::enzdes_out

-chemical:exclude_patches LowerDNA UpperDNA Cterm_amidation VirtualBB ShoveBB VirtualDNAPhosphate VirtualNTerm CTermConnect sc_orbitals pro_hydroxylated_case1 pro_hydroxylat^[d_case2 ser_phosphorylated thr_phosphorylated tyr_phosphorylated tyr_sulfated lys_dimethylated lys_monomethylated lys_trimethylated lys_acetylated glu_carboxylated cys_acetylated tyr_diiodinated N_acetylated C_methylamidated MethylatedProteinCterm

#-restore_talaris_behavior

###LIGAND SETUP###

#Only does torsions moving in ligand

-enzdes::minimize_all_ligand_torsions 10.0

#Moved ALL ligand torsions

-enzdes::detect_design_interface

-ligand::old_estat

#I/O

-overwrite

#-out:level 100

#Add hack_elec 0.25 to weights file

###ADDITIONAL PACKING SETUP###

-packing::extrachi_cutoff 1

-packing::ex1

-packing::ex2

-packing::ex1aro:level 6

-packing::ex2aro

-packing::extrachi_cutoff 1

-packing::use_input_sc

-packing::flip_HNQ

-packing::no_optH false

-packing::optH_MCA false

#-enzdes::favor_native_res 2

-enzdes::bb_min_allowed_dev 0.5

Parameters file (NAD^+^):

NAME NAD

IO_STRING NAD Z

TYPE LIGAND

AA UNK

ATOM O7 OH	X	-0.39

ATOM P1 Phos X	1.77

ATOM O1 OOC	X	-0.49

ATOM O3 OOC	X	-0.49

ATOM O12 OH	X	-0.39

ATOM C15 CH1	X	0.18

ATOM C11 CH1	X	0.18

ATOM C7 CH1	X	0.18

ATOM C4 CH1	X	0.18

ATOM C1 CH1	X	0.18

ATOM N7 Npro X	-0.10

ATOM C10 aroC X	0.15

ATOM N3 Nhis X	-0.26

ATOM C3 aroC X	0.15

ATOM N1 Nhis X	-0.26

ATOM C18 aroC X	0.15

ATOM C14 aroC X	0.15

ATOM N5 Nhis X	-0.26

ATOM C21 aroC X	0.15

ATOM N4 Nhis X	-0.26

ATOM O10 OH	X	-0.39

ATOM O4 OOC	X	-0.49

ATOM O8 OOC	X	-0.49

ATOM P2 Phos X	1.77

ATOM O2 OOC	X	-0.49

ATOM O6 OOC	X	-0.49

ATOM O13 OH	X	-0.39

ATOM C16 CH1	X	0.18

ATOM C12 CH1	X	0.18

ATOM C8 CH1	X	0.18

ATOM C5 CH1	X	0.18

ATOM C2 CH1	X	0.18

ATOM N2 Npro X	-0.10

ATOM C6 aroC X	0.15

ATOM C9 aroC X	0.15

ATOM C13 aroC X	0.15

ATOM C17 aroC X	0.15

ATOM C19 aroC X	0.15

ATOM C20 COO	X	0.89

ATOM N6 Nhis X	-0.26

ATOM O14 ONH2 X	-0.28

ATOM O11 OH	X	-0.39

ATOM O5 OOC	X	-0.49

ATOM O9 OOC	X	-0.49

BOND P1	O1

BOND P1	O3

BOND P1	O7

BOND P1	O12

BOND P2	O2

BOND P2	O6

BOND P2	O13

BOND C1	N7

BOND C2	N2

BOND N1	C3

BOND N2	C6

BOND N2	C19

BOND C3	N3

BOND C1	C4

BOND C4	O4

BOND C2	C5

BOND C5	O5

BOND C6	C9

BOND C4	C7

BOND C7	O8

BOND C5	C8

BOND C8	O9

BOND C9	C13

BOND C9	C20

BOND N3	C10

BOND P2	O7

BOND C7	C11

BOND C11 O10

BOND C8	C12

BOND C12 O11

BOND C13 C17

BOND C1	O10

BOND C2	O11

BOND C10 C14

BOND C14 C18

BOND C11 C15

BOND C12 C16

BOND C17 C19

BOND C15 O12

BOND C16 O13

BOND N1	C18

BOND C18 N4

BOND C20 N6

BOND C20 O14

BOND C14 N5

BOND N5	C21

BOND C10 N7

BOND C21 N7

CHI 1 P2	O7	P1	O1

CHI 2 O7	P1	O12 C15

CHI 3 O7	P2	O13 C16

CHI 4 C4	C1	N7	C10

CHI 5 C5	C2	N2	C6

CHI 6 C6	C9	C20 N6

CHI 7 P1	O7	P2	O2

CHI 8 O12 C15 C11 C7

CHI 9 O13 C16 C12 C8

CHI 10 P1	O12 C15 C11

CHI 11 P2	O13 C16 C12

NBR_ATOM O7

NBR_RADIUS 15.262751

ICOOR_INTERNAL	O7	0.000000	0.000000	0.000000	O7	P1	O1

ICOOR_INTERNAL	P1	0.000000 180.000000	1.542660	O7	P1	O1

ICOOR_INTERNAL	O1	0.000001	71.799358	1.483425	P1	O7	O1

ICOOR_INTERNAL	O3	128.835602	68.446545	1.484439	P1	O7	O1

ICOOR_INTERNAL	O12 113.355115	79.312579	1.607605	P1	O7	O3

ICOOR_INTERNAL	C15	90.808369	59.726547	1.438152	O12	P1	O7

ICOOR_INTERNAL	C11 124.522619	72.603621	1.507712	C15	O12	P1

ICOOR_INTERNAL	C7	-63.599121	66.616650	1.529059	C11	C15	O12

ICOOR_INTERNAL	C4 -114.139232	77.664591	1.524824	C7	C11	C15

ICOOR_INTERNAL	C1	-25.615437	78.033105	1.512552	C4	C7	C11

ICOOR_INTERNAL	N7	159.301237	65.581298	1.451857	C1	C4	C7

ICOOR_INTERNAL	C10 128.394631	54.068030	1.379551	N7	C1	C4

ICOOR_INTERNAL	N3	0.412496	52.500353	1.359237	C10	N7	C1

ICOOR_INTERNAL	C3 -179.715015	69.602108	1.314943	N3	C10	N7

ICOOR_INTERNAL	N1	0.195624	49.890783	1.371036	C3	N3	C10

ICOOR_INTERNAL	C18	-0.193855	62.555677	1.373392	N1	C3	N3

ICOOR_INTERNAL	C14	0.018997	62.508653	1.405869	C18	N1	C3

ICOOR_INTERNAL	N5	179.993873	48.596071	1.379811	C14	C18	N1

ICOOR_INTERNAL	C21 -179.755691	76.178960	1.326251	N5	C14	C18

ICOOR_INTERNAL	N4	179.858093	63.757573	1.315977	C18	N1	C14

ICOOR_INTERNAL	O10 -116.553710	77.045383	1.384677	C1	C4	N7

ICOOR_INTERNAL	O4 -122.937033	67.401015	1.445592	C4	C7	C1

ICOOR_INTERNAL	O8 -124.267796	70.690903	1.432167	C7	C11	C4

ICOOR_INTERNAL	P2	179.160793	48.208691	1.628696	O7	P1	O1

ICOOR_INTERNAL	O2	123.094195	74.331723	1.460404	P2	O7	P1

ICOOR_INTERNAL	O6 -122.431235	72.191363	1.489716	P2	O7	O2

ICOOR_INTERNAL	O13 -122.453996	80.675006	1.574622	P2	O7	O6

ICOOR_INTERNAL	C16 114.964724	62.717018	1.433805	O13	P2	O7

ICOOR_INTERNAL	C12 177.277878	71.813182	1.504974	C16	O13	P2

ICOOR_INTERNAL	C8	45.660747	61.443931	1.532631	C12	C16	O13

ICOOR_INTERNAL	C5	-92.661488	77.351149	1.522856	C8	C12	C16

ICOOR_INTERNAL	C2	-35.325058	78.297447	1.497911	C5	C8	C12

ICOOR_INTERNAL	N2	156.742316	66.914856	1.453748	C2	C5	C8

ICOOR_INTERNAL	C6	-65.483465	57.103063	1.408302	N2	C2	C5

ICOOR_INTERNAL	C9	178.822319	59.199989	1.488721	C6	N2	C2

ICOOR_INTERNAL	C13	2.241495	61.081207	1.511056	C9	C6	N2

ICOOR_INTERNAL	C17	-1.425405	66.965974	1.481730	C13	C9	C6

ICOOR_INTERNAL	C19	0.238385	54.300668	1.316589	C17	C13	C9

ICOOR_INTERNAL	C20 178.532536	62.237871	1.471225	C9	C6	C13

ICOOR_INTERNAL	N6	-0.436736	56.501852	1.334625	C20	C9	C6

ICOOR_INTERNAL	O14 179.935724	63.358536	1.240683	C20	C9	N6

ICOOR_INTERNAL	O11 -119.965361	74.196572	1.395439	C2	C5	N2

ICOOR_INTERNAL	O5 -119.545188	69.170240	1.440858	C5	C8	C2

ICOOR_INTERNAL	O9 -118.199430	70.697550	1.439752	C8	C12	C5

PDB_ROTAMERS NAD_conformers.pdb

Parameters file (NMN^+^):

NAME NMN

IO_STRING NMN Z

TYPE LIGAND

AA UNK

ATOM O5 OH	X	-0.41

ATOM C2 CH1	X	0.16

ATOM C1 CH1	X	0.16

ATOM O4 OH	X	-0.41

ATOM P1 Phos X	1.75

ATOM O1 OOC	X	-0.51

ATOM O2 OOC	X	-0.51

ATOM O3 OOC	X	-0.51

ATOM C3 CH1	X	0.16

ATOM O6 OOC	X	-0.51

ATOM C4 CH1	X	0.16

ATOM O7 OOC	X	-0.51

ATOM C5 CH1	X	0.16

ATOM N1 Npro X	-0.12

ATOM C6 aroC X	0.13

ATOM C7 aroC X	0.13

ATOM C8 COO	X	0.87

ATOM O8 ONH2 X	-0.30

ATOM N2 Nhis X	-0.28

ATOM C9 aroC X	0.13

ATOM C10 aroC X	0.13

ATOM C11 aroC X	0.13

BOND_TYPE O1	P1 2

BOND_TYPE P1	O2 1

BOND_TYPE P1	O3 1

BOND_TYPE P1	O4 1

BOND_TYPE O4	C1 1

BOND_TYPE C1	C2 1

BOND_TYPE C2	O5 1

BOND_TYPE C2	C3 1

BOND_TYPE O5	C5 1

BOND_TYPE C3	O6 1

BOND_TYPE C3	C4 1

BOND_TYPE C4	O7 1

BOND_TYPE C4	C5 1

BOND_TYPE C5	N1 1

BOND_TYPE N1	C6 2

BOND_TYPE N1	C11 1

BOND_TYPE C6	C7 1

BOND_TYPE C7	C8 1

BOND_TYPE C7	C9 2

BOND_TYPE C8	O8 2

BOND_TYPE C8	N2 1

BOND_TYPE C9	C10 1

BOND_TYPE C10 C11 2

CHI 1 C1	O4	P1	O1

CHI 2 C2	C1	O4	P1

CHI 3 O5	C2	C1	O4

CHI 4 C4	C5	N1	C6

CHI 5 C6	C7	C8	O8

NBR_ATOM O5

NBR_RADIUS 8.106243

ICOOR_INTERNAL	O5	0.000000	0.000000	0.000000	O5	C2	C1

ICOOR_INTERNAL	C2	0.000000 180.000000	1.505343	O5	C2	C1

ICOOR_INTERNAL	C1	0.000001	69.322004	1.371617	C2	O5	C1

ICOOR_INTERNAL	O4	127.026919	69.905303	1.430665	C1	C2	O5

ICOOR_INTERNAL	P1 -150.168026	58.832659	1.654853	O4	C1	C2

ICOOR_INTERNAL	O1	16.983450	69.507738	1.517317	P1	O4	C1

ICOOR_INTERNAL	O2	119.988765	70.757044	1.522187	P1	O4	O1

ICOOR_INTERNAL	O3	119.700537	70.583069	1.506982	P1	O4	O2

ICOOR_INTERNAL	C3 -121.087040	75.215482	1.607657	C2	O5	C1

ICOOR_INTERNAL	O6 -118.027897	69.788504	1.499394	C3	C2	O5

ICOOR_INTERNAL	C4	118.837141	74.015006	1.245008	C3	C2	O6

ICOOR_INTERNAL	O7 -139.868396	69.894037	1.482237	C4	C3	C2

ICOOR_INTERNAL	C5	119.230313	74.086588	1.658062	C4	C3	O7

ICOOR_INTERNAL	N1	159.309511	67.383726	1.461973	C5	C4	C3

ICOOR_INTERNAL	C6	36.107188	58.963419	1.349063	N1	C5	C4

ICOOR_INTERNAL	C7	179.777879	60.031912	1.384906	C6	N1	C5

ICOOR_INTERNAL	C8	179.820806	59.686733	1.535527	C7	C6	N1

ICOOR_INTERNAL	O8 -163.937877	59.936054	1.213332	C8	C7	C6

ICOOR_INTERNAL	N2 -179.863642	59.469442	1.440599	C8	C7	O8

ICOOR_INTERNAL	C9	179.870698	60.067994	1.351796	C7	C6	C8

ICOOR_INTERNAL	C10	0.027301	59.953999	1.350923	C9	C7	C6

ICOOR_INTERNAL	C11	0.083069	59.981151	1.370219	C10	C9	C7

PDB_ROTAMERS NMN_conformers.pdb

Constraint file (NAD^+^):

#NAD w/ T449

CST::BEGIN

TEMPLATE:: ATOM_MAP: 1	atom_name: O11

TEMPLATE:: ATOM_MAP: 1	residue3: NAD

TEMPLATE:: ATOM_MAP: 2	atom_name: OG1

TEMPLATE:: ATOM_MAP: 2	residue1: T

CONSTRAINT:: distanceAB: 3.7	0.2	50.0	0

\# CONSTRAINT:: angle_A: 180.0 5.0	500.0	360

\# CONSTRAINT:: angle_B: 120.0 5.0	500.0	360

\# CONSTRAINT:: torsion_B: 180.0 10.0 500.0	360

CST::END

#NAD w/ N643

CST::BEGIN

TEMPLATE:: ATOM_MAP: 1	atom_name: O14

TEMPLATE:: ATOM_MAP: 1	residue3: NAD

TEMPLATE:: ATOM_MAP: 2	atom_name: ND2

TEMPLATE:: ATOM_MAP: 2	residue1: N

CONSTRAINT:: distanceAB: 2.9	0.2	50.0	0

\# CONSTRAINT:: angle_A: 180.0 5.0	500.0	360

\# CONSTRAINT:: angle_B: 120.0 5.0	500.0	360

\# CONSTRAINT:: torsion_B: 180.0 10.0 500.0	360

CST::END

#NAD w/ I342

CST::BEGIN

TEMPLATE:: ATOM_MAP: 1	atom_name: O2

TEMPLATE:: ATOM_MAP: 1	residue3: NAD

TEMPLATE:: ATOM_MAP: 2	atom_name: N

TEMPLATE:: ATOM_MAP: 2	residue1: I

CONSTRAINT:: distanceAB: 4.0	0.2	50.0	0

\# CONSTRAINT:: angle_A: 180.0 5.0	500.0	360

\# CONSTRAINT:: angle_B: 120.0 5.0	500.0	360

\# CONSTRAINT:: torsion_B: 180.0 10.0 500.0	360

CST::END

#NAD w/ C479

CST::BEGIN

TEMPLATE:: ATOM_MAP: 1	atom_name: C13

TEMPLATE:: ATOM_MAP: 1	residue3: NAD

TEMPLATE:: ATOM_MAP: 2	atom_name: SG

TEMPLATE:: ATOM_MAP: 2	residue1: C

CONSTRAINT:: distanceAB: 3.5	0.3	50.0	0

\# CONSTRAINT:: angle_A: 180.0 5.0	500.0	360

\# CONSTRAINT:: angle_B: 120.0 5.0	500.0	360

\# CONSTRAINT:: torsion_B: 180.0 10.0 500.0	360

CST::END

#T426 w/ L428

CST::BEGIN

TEMPLATE:: ATOM_MAP: 1	atom_name: OG1

TEMPLATE:: ATOM_MAP: 1	residue1: T

TEMPLATE:: ATOM_MAP: 2	atom_name: N

TEMPLATE:: ATOM_MAP: 2	residue1: L

CONSTRAINT:: distanceAB: 3.2	0.2	50.0	0

\# CONSTRAINT:: angle_A: 180.0 5.0	500.0	360

\# CONSTRAINT:: angle_B: 120.0 5.0	500.0	360

\# CONSTRAINT:: torsion_B: 180.0 10.0 500.0	360

CST::END

#R341 w/ E644

CST::BEGIN

TEMPLATE:: ATOM_MAP: 1	atom_name: NE

TEMPLATE:: ATOM_MAP: 1	residue1: R

TEMPLATE:: ATOM_MAP: 2	atom_name: OE2

TEMPLATE:: ATOM_MAP: 2	residue1: E

CONSTRAINT:: distanceAB: 2.7	0.4	50.0	0

\# CONSTRAINT:: angle_A: 180.0 5.0	500.0	360

\# CONSTRAINT:: angle_B: 120.0 5.0	500.0	360

\# CONSTRAINT:: torsion_B: 180.0 10.0 500.0	360

CST::END

#T449 w/ Y647

CST::BEGIN

TEMPLATE:: ATOM_MAP: 1	atom_name: O

TEMPLATE:: ATOM_MAP: 1	residue1: T

TEMPLATE:: ATOM_MAP: 2	atom_name: OH

TEMPLATE:: ATOM_MAP: 2	residue1: Y

CONSTRAINT:: distanceAB: 2.9	0.2	50.0	0

\# CONSTRAINT:: angle_A: 180.0 5.0	500.0	360

\# CONSTRAINT:: angle_B: 120.0 5.0	500.0	360

\# CONSTRAINT:: torsion_B: 180.0 10.0 500.0	360

CST::END

#R340 w/ D363_1

CST::BEGIN

TEMPLATE:: ATOM_MAP: 1	atom_name: NH2

TEMPLATE:: ATOM_MAP: 1	residue1: R

TEMPLATE:: ATOM_MAP: 2	atom_name: OD1

TEMPLATE:: ATOM_MAP: 2	residue1: D

CONSTRAINT:: distanceAB: 3.0	0.3	100.0	0

\# CONSTRAINT:: angle_A: 180.0 5.0	500.0	360

\# CONSTRAINT:: angle_B: 120.0 5.0	500.0	360

\# CONSTRAINT:: torsion_B: 180.0 10.0 500.0	360

CST::END

#R340 w/ D363_2

CST::BEGIN

TEMPLATE:: ATOM_MAP: 1	atom_name: NE

TEMPLATE:: ATOM_MAP: 1	residue1: R

TEMPLATE:: ATOM_MAP: 2	atom_name: OD2

TEMPLATE:: ATOM_MAP: 2	residue1: D

CONSTRAINT:: distanceAB: 3.0	0.3	100.0	0

\# CONSTRAINT:: angle_A: 180.0 5.0	500.0	360

\# CONSTRAINT:: angle_B: 120.0 5.0	500.0	360

\# CONSTRAINT:: torsion_B: 180.0 10.0 500.0	360

CST::END

Constraint file (NMN^+^):

#NMN w/ T449

CST::BEGIN

TEMPLATE:: ATOM_MAP: 1	atom_name: O5

TEMPLATE:: ATOM_MAP: 1	residue3: NMN

TEMPLATE:: ATOM_MAP: 2	atom_name: OG1

TEMPLATE:: ATOM_MAP: 2	residue1: T

CONSTRAINT:: distanceAB: 3.7	0.2	50.0	0

\# CONSTRAINT:: angle_A: 180.0 5.0	500.0	360

\# CONSTRAINT:: angle_B: 120.0 5.0	500.0	360

\# CONSTRAINT:: torsion_B: 180.0 10.0 500.0	360

CST::END

#NMN w/ N643

CST::BEGIN

TEMPLATE:: ATOM_MAP: 1	atom_name: O8

TEMPLATE:: ATOM_MAP: 1	residue3: NMN

TEMPLATE:: ATOM_MAP: 2	atom_name: ND2

TEMPLATE:: ATOM_MAP: 2	residue1: N

CONSTRAINT:: distanceAB: 2.9	0.2	50.0	0

\# CONSTRAINT:: angle_A: 180.0 5.0	500.0	360

\# CONSTRAINT:: angle_B: 120.0 5.0	500.0	360

\# CONSTRAINT:: torsion_B: 180.0 10.0 500.0	360

CST::END

#NMN w/ I342

CST::BEGIN

TEMPLATE:: ATOM_MAP: 1	atom_name: O3

TEMPLATE:: ATOM_MAP: 1	residue3: NMN

TEMPLATE:: ATOM_MAP: 2	atom_name: N

TEMPLATE:: ATOM_MAP: 2	residue1: I

CONSTRAINT:: distanceAB: 4.5	0.2	50.0	0

\# CONSTRAINT:: angle_A: 180.0 5.0	500.0	360

\# CONSTRAINT:: angle_B: 120.0 5.0	500.0	360

\# CONSTRAINT:: torsion_B: 180.0 10.0 500.0	360

CST::END

#NMN w/ C479

CST::BEGIN

TEMPLATE:: ATOM_MAP: 1	atom_name: C9

TEMPLATE:: ATOM_MAP: 1	residue3: NMN

TEMPLATE:: ATOM_MAP: 2	atom_name: SG

TEMPLATE:: ATOM_MAP: 2	residue1: C

CONSTRAINT:: distanceAB: 3.5	0.3	50.0	0

\# CONSTRAINT:: angle_A: 180.0 5.0	500.0	360

\# CONSTRAINT:: angle_B: 120.0 5.0	500.0	360

\# CONSTRAINT:: torsion_B: 180.0 10.0 500.0	360

CST::END

#T426 w/ L428

CST::BEGIN

TEMPLATE:: ATOM_MAP: 1	atom_name: OG1

TEMPLATE:: ATOM_MAP: 1	residue1: T

TEMPLATE:: ATOM_MAP: 2	atom_name: N

TEMPLATE:: ATOM_MAP: 2	residue1: L

CONSTRAINT:: distanceAB: 3.2	0.2	50.0	0

\# CONSTRAINT:: angle_A: 180.0 5.0	500.0	360

\# CONSTRAINT:: angle_B: 120.0 5.0	500.0	360

\# CONSTRAINT:: torsion_B: 180.0 10.0 500.0	360

CST::END

#R341 w/ E644

CST::BEGIN

TEMPLATE:: ATOM_MAP: 1	atom_name: NE

TEMPLATE:: ATOM_MAP: 1	residue1: R

TEMPLATE:: ATOM_MAP: 2	atom_name: OE2

TEMPLATE:: ATOM_MAP: 2	residue1: E

CONSTRAINT:: distanceAB: 2.7	0.4	50.0	0

\# CONSTRAINT:: angle_A: 180.0 5.0	500.0	360

\# CONSTRAINT:: angle_B: 120.0 5.0	500.0	360

\# CONSTRAINT:: torsion_B: 180.0 10.0 500.0	360

CST::END

#T449 w/ Y647

CST::BEGIN

TEMPLATE:: ATOM_MAP: 1	atom_name: O

TEMPLATE:: ATOM_MAP: 1	residue1: T

TEMPLATE:: ATOM_MAP: 2	atom_name: OH

TEMPLATE:: ATOM_MAP: 2	residue1: Y

CONSTRAINT:: distanceAB: 2.9	0.2	50.0	0

\# CONSTRAINT:: angle_A: 180.0 5.0	500.0	360

\# CONSTRAINT:: angle_B: 120.0 5.0	500.0	360

\# CONSTRAINT:: torsion_B: 180.0 10.0 500.0	360

CST::END

Example of Rosetta script for docking (script for triple mutant shown here):

<ROSETTASCRIPTS>

<SCOREFXNS>

<ScoreFunction name="myscore" weights="ref2015_cst.wts"/>

</SCOREFXNS>

<SCORINGGRIDS ligand_chain="X" width="20.0">

<ClassicGrid grid_name="dock_grid" weight="1.0"/>

</SCORINGGRIDS>

<TASKOPERATIONS>

<DetectProteinLigandInterface name="edto" design="0" cut1="6.0" cut2="8.0" cut3="10.0" cut4="12.0"/>

<LimitAromaChi2 name="limchi2"/>

<SetCatalyticResPackBehavior name="catres" fix_catalytic_aa="0"/>

<ProteinLigandInterfaceUpweighter name="up" interface_weight="1.5" />

</TASKOPERATIONS>

<FILTERS>

<EnzScore name="allcst" score_type="cstE" scorefxn="myscore" whole_pose="1" energy_cutoff="10000000000000000000"/>

<LigInterfaceEnergy name="interfE" scorefxn="myscore" energy_cutoff="100000"/>

<ShapeComplementarity name="sc" min_sc="0.4" jump="1" />

</FILTERS>

<MOVERS>

<MutateResidue name="mutate1" target="%%target1%%" new_res="%%new_res1%%" />

<MutateResidue name="mutate2" target="%%target2%%" new_res="%%new_res2%%" />

<MutateResidue name="mutate3" target="%%target3%%" new_res="%%new_res3%%" />

<AddOrRemoveMatchCsts name="cstadd" cst_instruction="add_new"/>

<AddOrRemoveMatchCsts name="cstrem" cst_instruction="remove"/>

<PredesignPerturbMover name="predes1" trans_magnitude="1" rot_magnitude="20" dock_trials="1000"/>

<PredesignPerturbMover name="predes2" trans_magnitude="0.1" rot_magnitude="2.0" dock_trials="1000"/>

<EnzRepackMinimize name="desmin_nobb" design="0" repack_only="1" scorefxn_minimize="soft_rep" scorefxn_repack="soft_rep" minimize_rb="1" minimize_sc="1" minimize_bb="0" cycles="1" minimize_lig="1" min_in_stages="0" backrub="0" task_operations="edto,limchi2,up"/>

<EnzRepackMinimize name="desmin_wbb" design="0" repack_only="0" scorefxn_minimize="myscore" scorefxn_repack="soft_rep" minimize_rb="1" minimize_sc="1" minimize_bb="1" cycles="1" minimize_lig="1" min_in_stages="0" backrub="0" task_operations="edto,limchi2,up"/>

<EnzRepackMinimize name="cstopt" cst_opt="1" minimize_rb="1" minimize_sc="1" minimize_bb="0" cycles="1" min_in_stages="0" minimize_lig="1" />

<Transform name="trans1" chain="X" box_size="20" move_distance="2" angle="20" cycles="1000000" repeats="5" temperature="5"/>

<Transform name="trans2" chain="X" box_size="2" move_distance="0.2" angle="20" cycles="1000000" repeats="5" temperature="5"/>

<ParsedProtocol name="mutate">

<Add mover="mutate1" />

<Add mover="mutate2" />

<Add mover="mutate3" />

</ParsedProtocol>

<ParsedProtocol name="dock_des">

<Add mover="trans1"/>

<Add mover="trans2"/>

<Add mover="predes1"/>

<Add mover="predes2"/>

<Add mover="cstopt"/>

<Add mover_name="desmin_nobb"/>

<Add mover="desmin_wbb"/>

</ParsedProtocol>

<InterfaceScoreCalculator name="add_scores" chains="X" scorefxn="myscore" />

<GenericMonteCarlo name="multi_dock" mover_name="dock_des" filter_name="interfE" trials="8" />

</MOVERS>

<APPLY_TO_POSE>

</APPLY_TO_POSE>

<PROTOCOLS>

<Add mover_name="mutate" />

<Add mover_name="cstadd" />

<Add mover_name="multi_dock" />

<Add mover_name="add_scores" />

<Add filter_name="allcst" />

</PROTOCOLS>

</ROSETTASCRIPTS>
